# Inhibition of Asparagine Synthetase Effectively Retards Polycystic Kidney Disease Progression

**DOI:** 10.1101/2023.10.10.561720

**Authors:** Christine Podrini, Sara Clerici, Laura Tronci, Davide Stefanoni, Tamara Canu, Marco Chiaravalli, Daniel Spies, Ana S. H. Costa, Antonio Esposito, Angelo D’Alessandro, Christian Frezza, Angela Bachi, Alessandra Boletta

**Author notes:** Address Correspondence to: Alessandra Boletta, Division of Genetics and Cell Biology, San Raffaele Scientific Institute, Via Olgettina, 58, 20132, Milano, Italy. Equal Contributions.

## Abstract

Polycystic Kidney Disease (PKD) is a genetic disorder characterized by bilateral cyst formation. We showed that PKD cells and kidneys display metabolic alterations, including the Warburg effect and glutaminolysis, sustained *in vitro* by the enzyme asparagine synthetase (ASNS). Here, we used antisense oligonucleotides (ASO) against *Asns* in orthologous and slowly progressive PKD murine models and show that treatment leads to a drastic reduction of total kidney volume (measured by MRI) and a prominent rescue of renal function in the mouse. Mechanistically, the upregulation of an ATF4-ASNS axis in PKD is driven by the amino acid response (AAR) branch of the integrated stress response (ISR). Metabolic profiling of PKD or control kidneys treated with *Asns*-ASO or *Scr*-ASO revealed major changes in the mutants, several of which are rescued by *Asns* silencing *in vivo*. Indeed, ASNS drives glutamine-dependent *de novo* pyrimidine synthesis and proliferation in cystic epithelia. Notably, while several metabolic pathways were completely corrected by *Asns*-ASO, glycolysis was only partially restored. Accordingly, combining the glycolytic inhibitor 2DG with *Asns*-ASO further improved efficacy. Our studies identify a new therapeutic target and novel metabolic vulnerabilities in PKD.

## Introduction

Autosomal Dominant Polycystic Kidney Disease (ADPKD) is one of the most common monogenic inheritable disorders. It is caused by loss of function in either *PKD1* (85% of the cases) or *PKD2* (15% of the cases) genes, which encode for polycystins, PC-1 and PC-2 respectively (Torres *et al*, 2007; Harris & Torres, 2014; Ong & Harris, 2015). The inheritance of one mutated allele is not sufficient for the establishment of the pathology, thus it has been proposed that a second hit causing loss of heterozygosity is needed to drop polycystins’ complex functional activity under a critical threshold (Qian *et al*, 1996; Watnick *et al*, 2000; Leeuwen *et al*, 2004; Hopp *et al*, 2012). The main clinical manifestation of the disease is bilateral formation and expansion of numerous cysts, that progressively compress and compromise the plasticity and function of the kidney. Secondary implications include cyst development in the liver and pancreas (Torres *et al*, 2007; Harris & Torres, 2014; Ong & Harris, 2015). In all these tissues, cysts are focal, clonal, and fluid-filled structures outpunching from the tubular epithelium (Qian *et al*, 1996). These cystic structures progressively increase in dimension and number during the lifespan of the patient (Harris & Torres, 2014; Bergmann *et al*, 2018). Furthermore, cardiovascular complications, hypertension, and aneurism are important features of the disease (Bergmann *et al*, 2018). Notably, molecular mechanisms driving the cystogenic process are still not completely uncovered. However, different pathways are known to be dysregulated during the cyst expansion process, and likely important to determine declining renal function.

Only one compound has been developed up to registration for this disease, based on an antagonist of the Vasopressin type II receptor (called Tolvaptan) that hampers cAMP production in the collecting ducts, driving increased proliferation, as observed in preclinical models and in human samples (Torres *et al*, 2017). While the development of Tolvaptan proved the feasibility of completing all the regulatory pathway for this complicated and life-long disorder, the molecule has limited efficacy, is poorly tolerated, and presents with quite important side effects such as rare, but severe, liver toxicity (Torres *et al*, 2017; Watkins *et al*, 2015). Thus, the development of new therapeutic approaches for ADPKD is mandatory.

Our laboratory, among others, has contributed to defining ADPKD as a metabolic disorder, describing how the metabolic rewiring supports the required increase in proliferation observed during cystogenesis, and opening interesting opportunities for therapy (Rowe *et al*, 2013; Chiaravalli *et al*, 2016; Podrini *et al*, 2018, 2020). Metabolic reprogramming might be a particularly appealing dysfunction to be treated in the disease, as it is quite prominent and sustains the energetic requirements of proliferation. For instance, targeting the increased glycolysis using 2-deoxy-D-glucose (2-DG) treatment resulted in a prominent improvement of disease progression in preclinical animal models of the disease (Rowe *et al*, 2013; Chiaravalli *et al*, 2016; Riwanto *et al*, 2016; Nikonova *et al*, 2018; Lian *et al*, 2019). Abnormal cystic growth relies on the utilization of aerobic glycolysis, in a way resembling the Warburg effect observed in cancer, whereby cells lacking the polycystins become dependent on glucose for energy production (Rowe *et al*, 2013). Using metabolic tracing studies relying on heavy isotopologues we have demonstrated that glutaminolysis is increased, in what we have described to be a compensatory mechanism able to fuel the TCA cycle (Podrini *et al*, 2018). Small amounts of this glutamine are utilized on the oxidative side of the TCA cycle allowing for the maintenance of mitochondrial membrane potential, even if glutamine utilization does not compensate for the severely reduced OXPHOS levels in *Pkd1^-/-^* cells (Menezes *et al*, 2016; Padovano *et al*, 2017; Podrini *et al*, 2018). Quite abundant levels of glutamine-derived a-KG are instead diverted towards reductive carboxylation, to generate citrate which provides abundant acetyl-CoA levels required to fuel fatty acids biosynthesis, sustaining membrane synthesis, and supporting proliferation, at least *in vitro* (Podrini *et al*, 2018).

Thus, we and others have concomitantly highlighted cysts’ dependence on glutamine utilization (Flowers *et al*, 2018; Podrini *et al*, 2018; Soomro *et al*, 2018), and we have further demonstrated that asparagine synthetase (ASNS) is a key player in the proficient utilization of glutamine to fuel TCA cycle and to cope with the increased energetic demand (Podrini *et al*, 2018). Indeed, silencing of *Asns in vitro* drastically impaired the glutamine utilization of *Pkd1^-/-^* cells leading to compromised survival and proliferation (Podrini *et al*, 2018). Of note, an increase in circulating asparagine has subsequently been reported in children and young adults with ADPKD (Baliga *et al*, 2021), supporting the likely increased expression or activity of this enzyme in PKD patients. Similarly, different types of solid tumors as well as leukemia have been associated with high expression levels of the enzyme ASNS, which can promote proliferation, metastasis, and chemoresistance (Chiu *et al*, 2020).

Here, we aimed to investigate the potential role of ASNS as a novel target for therapy acting on a metabolic vulnerability of ADPKD. We show that targeting *Asns* with antisense oligonucleotides (ASOs) in a slowly progressive orthologous model of PKD results in strong and remarkable disease improvement. Moreover, we identified general control nonderepressible 2 (GCN2) as a major regulator of ASNS increased expression in this disease, mediated by an activation of the amino acid response (AAR) branch of the integrated stress response (ISR). Importantly, metabolomic profiling confirmed the increased asparagine levels in PKD kidneys and the concomitant upregulation of glycolysis and glutaminolysis. Notably, *Asns*-directed ASOs rescue some of the metabolic alterations observed in PKD, without affecting glycolysis. In line with this, combining *Asns* inhibition with 2DG provides a further delay in disease progression. Finally, in the attempt to better understand the mechanism of action of ASNS upregulation in ADPKD, we found that its inhibition blunted the Carbamoyl-phosphate synthetase 2, Aspartate transcarbamoylase, and Dihydroorotase (CAD)- dependent *de novo* pyrimidine synthesis pathway. Indeed, analysis of ^13^C_5_-glutamine tracing using cells lacking *Pkd1* revealed an upregulation of this metabolic pathway.

Our data collectively demonstrate that targeting ASNS is a new and effective therapeutic approach which deserves to be further exploited.

## Results

### ASNS is upregulated in ADPKD

Asparagine synthetase (ASNS) is a transamidase that catalyzes the synthesis of asparagine (Asn) from aspartate (Asp) by deamidating glutamine (Gln) to form glutamate (Glu), through an ATP-dependent reaction (Lomelino *et al*, 2017). Despite being ubiquitous, ASNS is generally expressed at low levels in organs other than the exocrine pancreas (Milman & Cooney, 1974; Uhlén *et al*, 2015). However, the enzyme is tightly regulated as a biosensor of cell stress stimuli, and it was found to be induced in pathological conditions including ADPKD (Lomelino *et al*, 2017; Podrini *et al*, 2018). Importantly, an increase in ASNS metabolic activity has been inferred in children and young adult ADPKD patients based on the increased levels of circulating asparagine (Baliga *et al*, 2021). Thus, we investigated whether ASNS was indeed upregulated using both *in vitro* and *in vivo* PKD models. Consistent with our previously reported transcriptional upregulation (Podrini *et al*, 2018), we found increased ASNS protein in Mouse Embryonic Fibroblasts (MEF) knockout for the *Pkd1* gene compared to controls, in basal conditions. Interestingly, ASNS expression was further increased in *Pkd1^-/-^* MEFs upon glucose deprivation both at the mRNA (**Fig 1A**) and protein level (**Fig 1B**), in line with the idea that ASNS mediates the glutamine compensatory utilization and with previous evidence in cancer (Chiu *et al*, 2020). Moreover, we observed increased ASNS protein expression in cystic *KspCre;Pkd1^ΔC/flox^*kidneys, which develop early and severe PKD at post-natal day 4 (P4) (**Fig 1C**) compared to relative controls. Further, analysis of published data from different PKD animal models, *Pkd1^v/v^* mice at P10 (**Fig 1D**; (Podrini *et al*, 2018)) and *Pkd1^RC/RC^* mice at P12 (**Fig 1E**; (Olson *et al*, 2019)) showed an increased *Asns* transcript in these models as well. We confirmed the upregulation of *ASNS* in ADPKD human samples microarray (**Fig 1F**; (Song *et al*, 2009)), as previously observed (Podrini *et al*, 2018). Finally, analysis of previously published datasets of single nucleus RNA-sequencing of ADPKD patients (Muto *et al*, 2021) confirmed *ASNS* increase in clusters of proximal and distal tubules from cystic tissues compared to controls (**Fig 1G**). Together, these data support the evidence that ASNS is strongly upregulated in multiple cellular and animal models as well as in human ADPKD samples.

**Figure 1.**
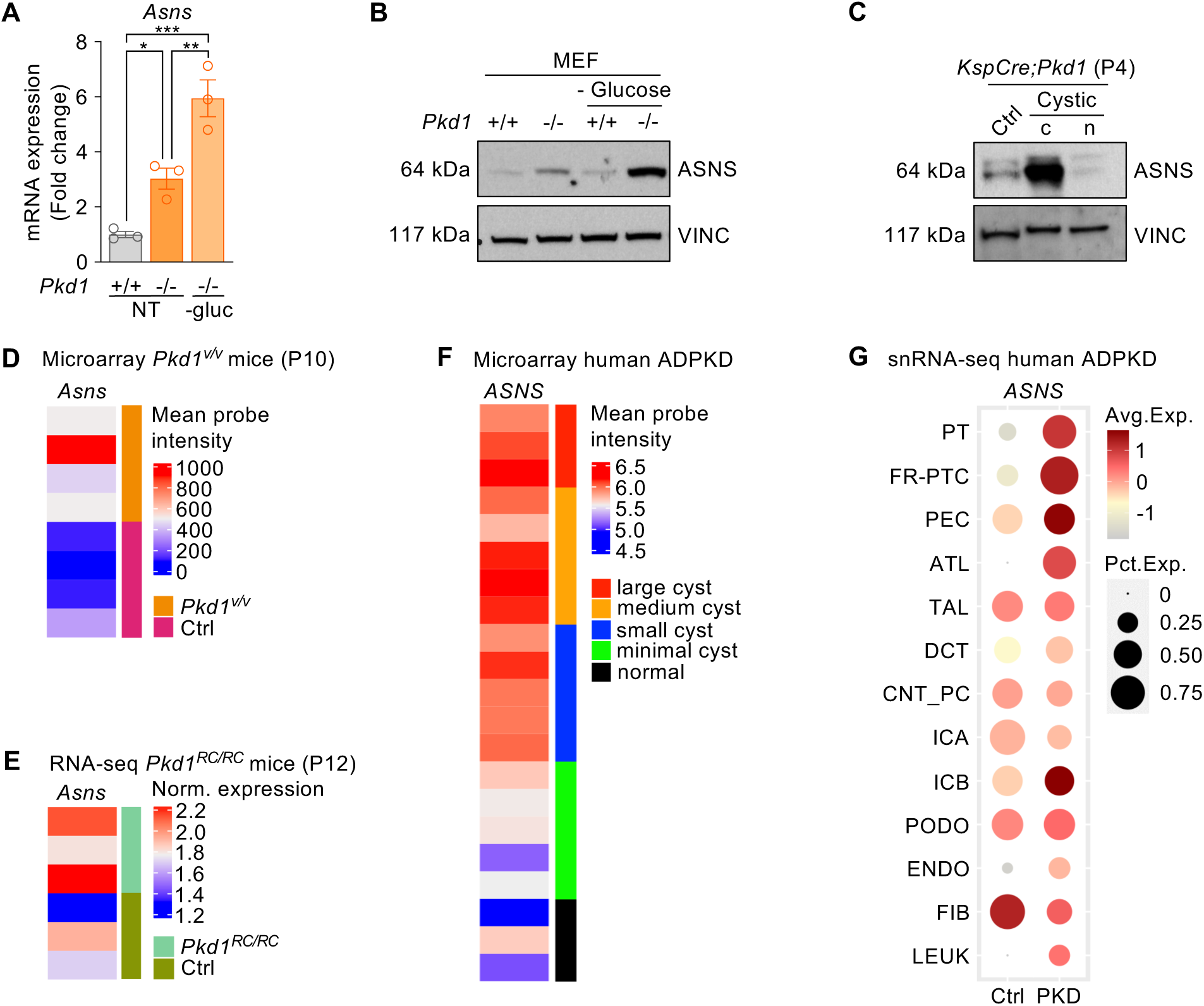
ASNS is upregulated in PKD models and in human ADPKD. A. *Asns* mRNA expression in *Pdk1^-/-^* and control MEF, cultured in full medium or under glucose deprivation. Representative of n=3 independent experiments. B. ASNS expression in *Pdk1^-/-^* and control MEF, cultured in full medium or under glucose deprivation. Representative of n=3 independent experiments. C. ASNS expression in *KspCre;Pkd1^ΔC/flox^* (cystic) and relative control kidneys at P4. c cytoplasmatic fraction, n nuclear fraction. D. *Asns* expression in P10 *Pkd1^v/v^* mice microarray. E. *Asns* expression in P12 *Pkd1^RC/RC^*mice RNA-seq. F. *ASNS* expression in ADPKD human samples microarrays. G. Dot plot of snRNA-seq dataset showing *ASNS* expression in clusters identified in human ADPKD cystic and normal kidney tissues. PT proximal tubule, FR-PTC failed-repair proximal tubular cells, PEC parietal epithelial cells, TAL thick ascending limb of Henle’s loop, DCT distal convoluted tubule, CNT_PC connecting tubule and principal cells, ICA Type A intercalated cells, ICB Type B intercalated cells, PODO podocytes, ENDO endothelial cells, FIB fibroblasts, LEUK leukocytes. Data information: in A data are shown as mean ± SD. One-way ANOVA. *p<0.05; **p<0.01; ***p<0.001.

### Targeting *Asns* ameliorates key features of PKD progression

We have previously demonstrated that ASNS is a central player in the metabolic rewiring occurring in PKD, being the preferential enzyme for glutamine usage in *Pkd1* mutant cells. Indeed, its silencing impaired proliferation and enhanced *Pkd1^-/-^* cell apoptosis (Podrini *et al*, 2018). Thus, we investigated the effect of targeting ASNS *in vivo* in an orthologous and slowly progressive PKD mouse model carrying a Tamoxifen-inducible Cre recombinase (B6.Cg-Tg(CAG-cre/Esr1*)5Amc/J) enabling the inactivation at different time points (*Tam-Cre;Pkd1^ΔC/flox^*) (Piontek *et al*, 2007; Wodarczyk *et al*, 2009). Given the lack of specific inhibitors of the enzyme, mice were treated with anti-sense oligonucleotides against *Asns* (*Asns*-ASO) or scramble controls (*Scr*-ASO). A pilot study was conducted to assess the efficacy of *Asns* silencing in a small cohort of mice carrying *Pkd1* gene inactivation upon Tamoxifen (250 mg/kg) injection at P25 and developing a cystic phenotype within the next 70 days following a previously established protocol (Chiaravalli *et al*, 2016) (**Fig EV1A**). Intra-peritoneal (i.p.) injection of ASOs (50 mg/kg) weekly for 5 weeks and every other week till the end of the study was sufficient to substantially reduce the *Asns* transcript in kidney tissues (**Fig EV1B**), indicating that this dosage should be sufficient to silence the transcript. Of interest, we observed a trend of reduction in kidney volume (**Fig EV1C**), kidney over body weight (**Fig EV1D**) and Blood Urea Nitrogen levels (BUN, **Fig EV1E**), despite the low numerosity of this pilot cohort of mice. Histological analysis of transversal kidney sections showed a reduction in the percentage of cystic area in *Asns*-ASO treated group, compared to *Scr* one (**Fig EV1F**). Given the high variability of the model and the relatively low number of samples in this pilot study, the data pointed to a possible quite robust effect of improvement in disease progression upon silencing of *Asns*, and we set out to test the hypothesis using a more robust experimental design.

We designed a study that could rely on a longer treatment using the same mouse model (*Tam- Cre;Pkd1^ΔC/flox^*), in which the *Pkd1* gene was inactivated by Tamoxifen (250 mg/kg) injection at postnatal day 45 (P45) leading to the development of a slowly progressive disease that enabled treatment for over four months (up to P160, (Leeuwen *et al*, 2007)). *Asns*- and *Scr*-ASOs (50 mg/kg) were i.p. injected weekly up to P100, and every other week thereafter until the end of the study. This approach allowed to analyze over time different parameters linked to pathology progression as indicated in the experimental design (**Fig 2A**). At sacrifice, we confirmed that ASNS is induced in cystic kidneys at P160 and almost completely abrogated by *Asns*-ASO administration as compared to matching *Scr-*ASO treated animals (**Fig 2B**). Similar results were obtained when looking at the protein level, though some higher variability could be appreciated (**Fig 2C**). Magnetic resonance imaging (MRI) was applied to follow the progressive increase in kidney volume of the cystic mice after enrollment in the study. Interestingly, MRI scans along time (**Fig 2D**) and the relative quantification (**Fig 2E**) highlighted a strong reduction in kidney volume at P130 in the *Asns-*ASO treated group compared to the *Scr*-ASO one. Notably, *Asns*-ASO treatment completely rescued the renal function impairment, evaluated as blood urea nitrogen (BUN) (**Fig 2F**), and partially rescued the increased creatinine levels (**Fig 2G**). At the endpoint, *Asns*-ASO-treated animals showed reduced kidneys/body weight (**Fig 2I**) compared to cystic *Scr*-treated ones, as clearly appreciable through representative images of an average experiment (**Fig 2H**). Consistently, the histological analysis of kidney sections highlighted a reduction in the percentage cystic area (**Fig 2J**) in *Asns*-ASO group compared to *Scr* one, with some of the tissues analyzed showing an almost complete restoration of healthy tubules and parenchyma (**Fig 2K** and **Appendix Fig S1**). This study suggests that silencing *Asns* is a valuable approach to ameliorating key features of PKD and delaying disease progression.

**Figure 2.**
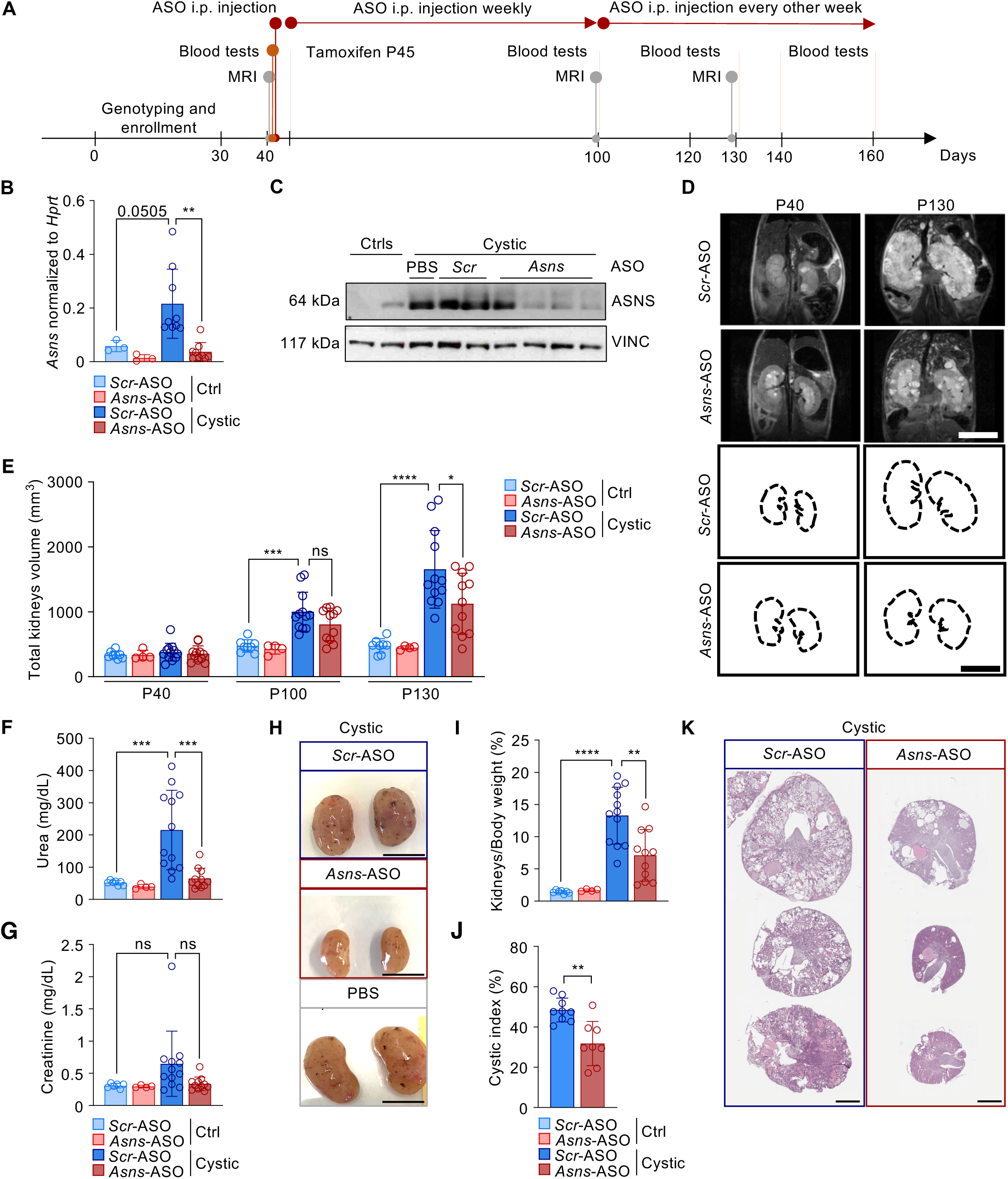
Targeting ASNS ameliorates PKD phenotype. A. Experimental design of *Asns*-ASO-treatment study in *Tam-Cre;Pkd1^ΔC/flox^* model (n=8 ctrls *Scr*-ASO; n=4 ctrls *Asns*-ASO; n=12 cystic *Scr*-ASO; n=11 cystic *Asns*-ASO). B. *Asns* mRNA expression in cystic kidneys compared to controls at P160, treated with either *Scr*-ASO or *Asns*-ASO. C. ASNS protein expression in cystic kidneys, treated with either PBS, *Scr*-ASO or *Asns*-ASO, compared to controls. D. MRI representative images of cystic kidneys treated with *Scr*-ASO, *Asns*-ASO. Images were acquired at P40 (before tamoxifen induction) and at P130. E. Total kidneys volume of cystic and control mice treated with *Scr*-ASO or *Asns*-ASO, calculated at P40, P100 and P130. F. BUN concentration in cystic and control mice treated with *Scr*-ASO or *Asns*-ASO, measured at P160. G. Creatinine concentration in cystic and control mice treated with *Scr*-ASO or *Asns*-ASO, measured at P160. H. Representative images of cystic kidneys harvested at P160 from mice treated with *Scr*-ASO, *Asns*-ASO or PBS. Scale bar (1 cm). I. Percentage of kidney weight normalized on total body weight at P160 of mice treated with *Scr*-ASO or *Asns*-ASO. J. Quantification of the cystic area percentage of the total kidney area measured in sections of cystic *Scr*-ASO or *Asns*-ASO groups. K. Representative images of H&E stained cross sections of P160 cystic kidneys, treated with *Scr*-ASO or *Asns*-ASO. Scale bar (2 mm). Data information: in B, E-G, I, data are shown as mean ± SD. One-way ANOVA. ns non-significant; *p<0.05; **p<0.01; ***p<0.001; ****p<0.0001. In J data are shown as mean ± SD. Student’s unpaired two-tailed t-test. **p<0.01.

### The GCN2-dependent branch of the integrated stress response drives ASNS upregulation in PKD

Given the primary role of ASNS in supporting ADPKD progression, we investigated the mechanism that underlies its upregulation. ASNS expression and activity in humans are highly prompted by cell stress. Indeed, this enzyme is transcriptionally regulated by activating transcription factor 4 (ATF4), which sits at the crossroad of two main branches of the integrated stress response (ISR) (**Fig 3A**), namely the amino acids response (AAR) and the unfolded protein response (UPR). Notably, we found that ATF4 is upregulated in all PKD models described above, including *KspCre;Pkd1^ΔC/flox^* kidneys at P4 (**Fig 3B**) and P45-tamoxifen-induced *Tam-Cre;Pkd1^ΔC/flox^* kidneys at P160 (**Fig 3C**). Of interest, the ATF4-ASNS axis is known to be activated in response to ER stress by PERK-dependent activation of unfolded protein response (UPR), a pathway that has been shown not to be de-regulated in PKD (Fedeles *et al*, 2015; Roy *et al*, 2023), but also during an imbalance in amino acids availability in cells and tissues via activation of the General Control Nonderepressible-2- kinase (GCN2) within the so called amino acid response (AAR) that converges downstream with the PERK pathway on eIF2alpha which translationally regulates ATF4 levels (Lomelino *et al*, 2017) (**Fig 3A**). This process is mimicked by the limitation of essential amino acids (EAAs) in cell culture (Jin *et al*, 2021), whereby GCN2 acts as a metabolic sensor for the depletion of amino acids, via the binding of uncharged tRNAs (Kanno *et al*, 2020). Our previous metabolomic analysis of *KspCre;Pkd1^ΔC/flox^* cystic kidneys in newborn mice highlighted an impairment in the aminoacyl-tRNA pathway as well as an imbalance in amino acid biosynthesis (Podrini *et al*, 2018), suggesting that the amino acid response (AAR) could be the driver for ASNS upregulation in PKD. Thus, we investigated if ASNS expression is regulated by the GCN2-dependent branch of the integrated stress response (ISR). Indeed, we found a trend of upregulation of GCN2 transcript in *Pkd1^-/-^* MEF cells (**Fig 3D**) along with an upregulation of the phosphorylation (**Fig 3F**) levels, which was paralleled by ATF4 and ASNS increase. Furthermore, *Eif2ak4* (the GCN2 gene) silencing completely abrogated ATF4 upregulation and ASNS induction (**Fig 3E-F**), indicating that this kinase is essential in the process. Collectively, these data show that *Pkd1* loss leads to upregulation of the amino acid response (AAR) which ultimately is responsible for the upregulation of the enzyme ASNS.

**Figure 3.**
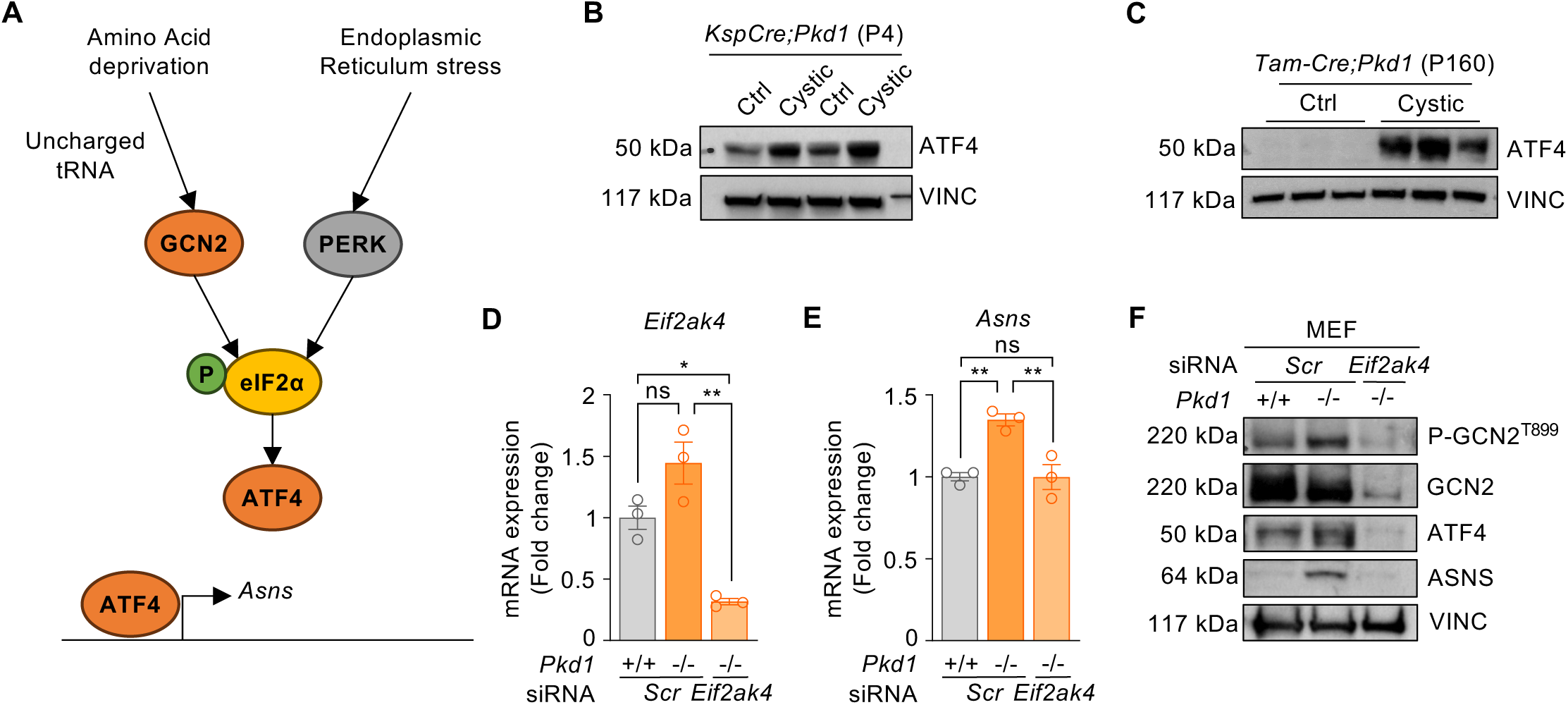
ASNS is upregulated via GCN2-dependent activation of ATF4. A. Schematic representation of the integrated stress response-dependent transcription of *Asns*. B. ATF4 protein expression in *KspCre;Pkd1^ΔC/flox^* cystic kidneys and relative controls at P4. C. ATF4 protein expression in *Scr*-ASO-treated *Tam-Cre;Pkd1^ΔC/flox^* mice and relative controls at P160. D. *Eif2ak4* expression in *Pkd1^-/-^* and control MEF cells upon silencing (representative of n=2). E. *Asns* expression in *Pkd1^-/-^* and control MEF cells upon *Eif2ak4* silencing (representative of n=3) F. P-GCN2, ATF4 and ASNS protein expression in *Pdk1^-/-^* and control MEF ± silencing of *Eif2ak4* (n=1). Data information: in D-E data are shown as mean ± SD. One-way ANOVA. Ns not-significant; *p<0.05; **p<0.01.

### *Asns*-ASO treatment rescues several metabolic pathways in PKD

In our previous studies, we identified ASNS as a key player in glutaminolysis in PKD. Therefore, we set out to determine whether *Asns*-ASO treatment impacts on PKD progression in our *in vivo* model by affecting key metabolic processes deregulated in the disease. To assess this, we performed targeted metabolomics by liquid-chromatography mass-spectrometry (LC-MS) on cystic and control kidneys treated with *Asns*-ASO versus *Scr*-ASO. To this end, and to facilitate the identification of profound changes in metabolism, we selected the five cystic *Scr*-ASO-treated mice with the most exacerbated PKD phenotype, and the five cystic *Asns*-ASO-treated animals that presented the most striking improvement, as assessed by MRI scans at P130 (**Fig 4A**). Profiling using LC-MS identified 162 metabolites, out of a total of 265, that were changing significantly (p<0.05) between conditions. Notably, principal component analysis (PCA) revealed a clear clustering of controls *Scr*-ASO and *Asns*-ASO indicating a minimal impact of this treatment on WT kidneys, in line with the low levels of expression of this enzyme in basal conditions. Of note, *Scr*-ASO cystic kidneys clustered together and separated quite robustly from the controls. On the other hand, cystic kidneys treated with *Asns*-ASO clustered together and almost completely overlapped with the controls, indicating that the treatment rescued the metabolic derangement in PKD kidneys almost to completion (**Fig 4B**). The dendrogram (**Fig EV2A**) and hierarchical clustering analysis (HCA) further confirmed this clustering of the samples (**Fig EV2B**). Furthermore, heatmap representation clearly showed that targeting ASNS affects the metabolic processes altered in PKD cystic kidneys by rescuing 53.70% of the 162 metabolites altered in *Scr*-ASO ones (**Fig EV2B**). We first confirmed that silencing of *Asns* significantly reduced asparagine levels, which were increased in cystic *Scr*-ASO kidneys, in line with the functional role of this enzyme. The substrates (Aspartate and Glutamine), as well as the second product of the reaction (Glutamate), were consistently changing in *Scr*-ASO versus *Asns*-ASO (**Fig 4C**). Moreover, *Asns*-ASO treatment reduced Citrate and α-KG levels, which could be explained by a decrease in reductive carboxylation, in line with our previous observation in *Pkd1^-/-^* MEF cells upon silencing of *Asns* (Podrini *et al*, 2018), and resulting in glutamate accumulation (**Fig 4C**). According to previous studies (Rowe *et al*, 2013), glycolysis was upregulated in cystic kidneys, in line with the robust Warburg effect observed in this disease, but *Asns*-ASO only partially reduced glucose and lactate levels (**Fig 4C**). Indeed, pathway enrichment analysis based on KEGG database identified 11 upregulated (dark red) and 5 downregulated (dark blue) metabolic pathways significantly changed in cystic *Scr*-ASO kidneys compared to the other groups. Notably, 10 out of the 16 pathways were completely rescued by *Asns*-ASO treatment, as they were no longer significantly different compared to control groups (**Fig 4D**). Interestingly, various pathways were related to amino acid metabolism and aminoacyl-tRNA biosynthesis (**Fig 4D**) and rescued by the silencing of *Asns*. Indeed, we observed an imbalance of amino acids in cystic kidneys, involving some essential aa, which was corrected by *Asns*-ASO treatment (**Fig EV2C**). This possibly supports the observation that *Asns* upregulation is mediated by the AAR branch of the ISR in cystic kidneys, as described in **Fig 3**. Altogether, these results corroborate the central role of ASNS in the metabolic rewiring occurring in PKD, highlighting the therapeutic potential of its inhibition.

**Figure 4.**
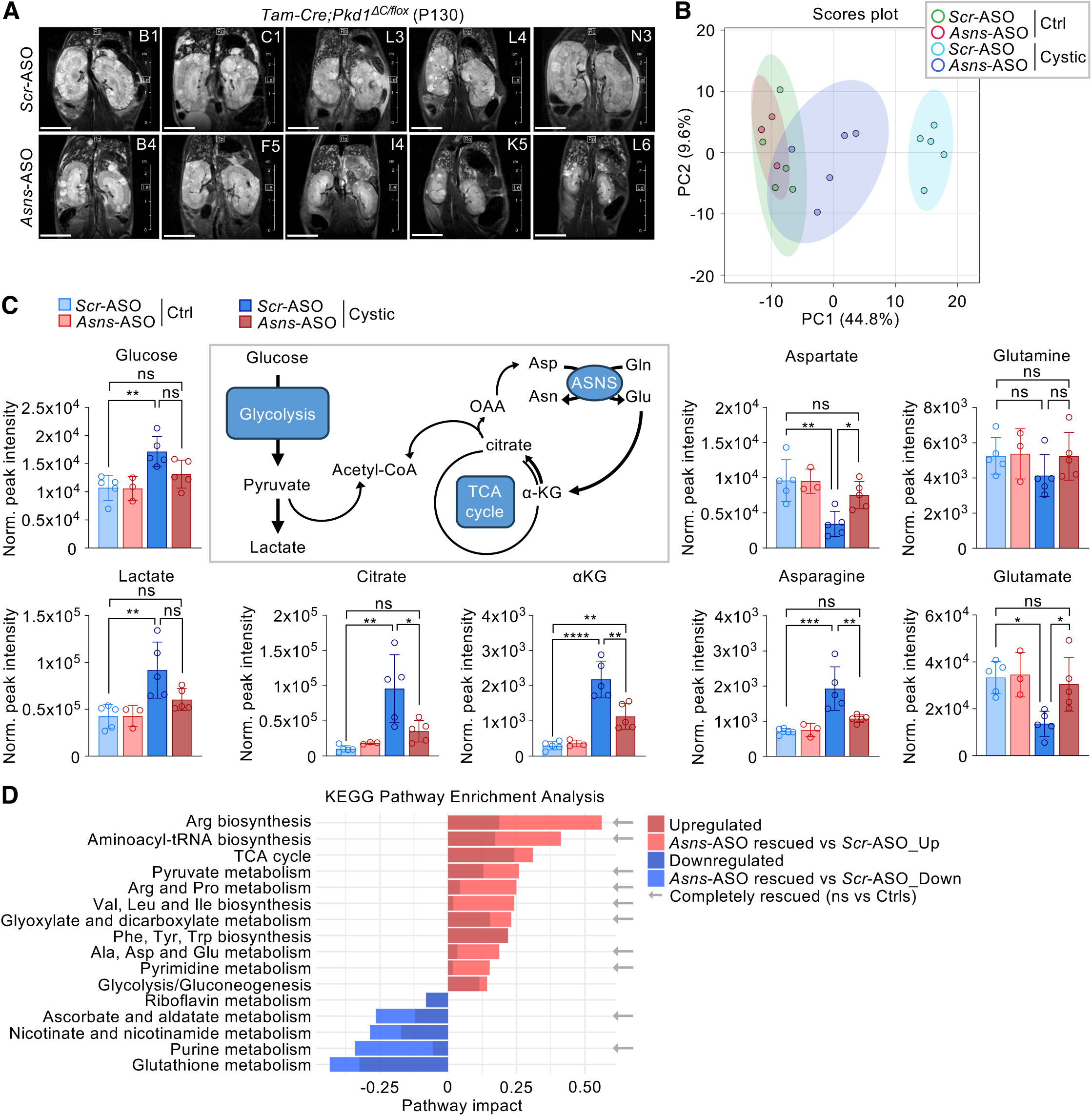
*Asns*-ASO treatment rescues the metabolic reprogramming occurring in PKD. A. MRI scan of P130 cystic *Tam-Cre;Pkd1^ΔC/flox^*kidneys, treated with ASOs, included in the metabolomic analysis. B. Principal Component Analysis (PCA) of targeted metabolomics of *Tam-Cre;Pkd1^ΔC/flox^* cystic kidneys and relative controls, treated with *Scr*- or *Asns*-ASO (n=5 ctrl *Scr*-ASO; n=3 ctrl *Asns*-ASO; n=5 cystic *Scr*-ASO; n=5 cystic *Asns*-ASO). C. Schematic representation and analysis of metabolites involved in ASNS-dependent fueling of TCA cycle (reductive carboxylation) and glycolysis. D. Pathway Enrichment Analysis of Up- and Down-regulated pathways (Cystic *Scr*-ASO respect to the other groups), and rescue of *Asns*-ASO treatment compared to *Scr*-ASO group. Pathways non-statistically (ns) different from Ctrl group upon *Asns*-ASO treatment are indicated as completely rescued. Data information: In C data are shown as mean ± SD. One-way ANOVA, corrected with Tukey’s multiple comparisons. ns non-significant; *p<0.05; **p<0.01; ****p<0.0001. In D statistical significance of Pathway Enrichment and Impact was evaluated with Hypergeometric Test with Relative-betweenness Centrality, based on KEGG Database with p<0.05, FDR<0.1.

### ASNS sustains *de novo* pyrimidine synthesis in PKD

One of the central mechanisms through which ASNS sustains proliferation in various cancerous conditions is the increased *de novo* pyrimidine biosynthesis (Sullivan *et al*, 2015; Krall *et al*, 2016, 2021). In line with this, our data showed that this pathway was strongly upregulated in PKD kidneys, potentially supporting proliferation during the cystic process, and was completely rescued by *Asns* targeting (**Fig 4D**). Indeed, looking at the individual metabolites identified in the LC-MS profiling of ASO-treated kidneys revealed that all three intermediate products (Carbamoyl phosphate, N-Carbamoyl aspartate, Dihydroorotate) of the multifunctional enzyme CAD (Carbamoyl-phosphate synthetase 2, Aspartate transcarbamoylase and Dihydroorotase) were upregulated in cystic kidneys and rescued by *Asns*-ASO (**Fig 5A**). Consistently, the intermediates of the following step of nucleotide biosynthesis (Orotate, Orotidine-5’-phosphate) were increased in diseased kidneys and responsive to ASO treatment (**Fig 5A**). Furthermore, the reduction of the substrates UMP (Uridine-5’- monophosphate) and PRPP (5-phospho-α-D-ribose-1-diphosphate) supports the idea of a possible increased consumption for pyrimidine biosynthesis (Thymine, Cytosine, Uracil), and it is also rescued by *Asns* targeting (**Fig 5A**). Given the major effect of *Asns* targeting on pyrimidine biosynthesis, which is required to support proliferation, and the evidence that silencing of *Asns* impairs the proliferation of *Pkd1^-/-^* cells (Podrini *et al*, 2018), we tested whether *Asns*-ASO treatment could hamper cyst expansion in the *Tam-Cre;Pkd1^ΔC/flox^* kidneys acting on the proliferative potential of epithelial mutant cells. Thus, we quantified the percentage of Ki67-positive nuclei in the epithelium lining the cysts of ASO-treated kidneys. We observed a reduction in the proliferation of *Asns*-ASO treated kidneys of the pilot study at P94, compared to *Scr*-treated samples (**Fig 5B**).

**Figure 5.**
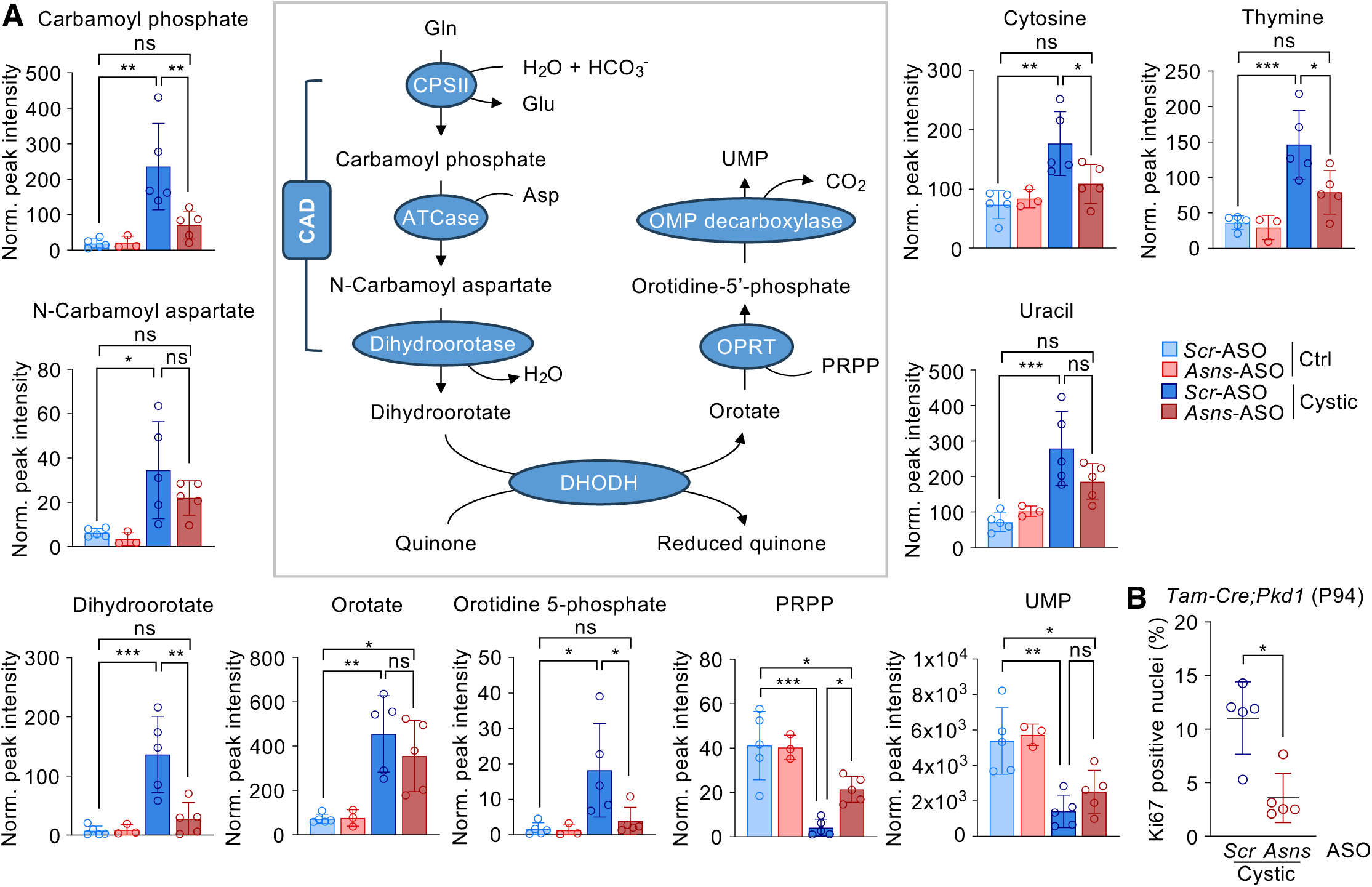
Targeting ASNS hampers CAD-dependent pyrimidine biosynthesis in PKD model. A. Schematic representation CAD-dependent *de novo* pyrimidine biosynthesis pathway and targeted metabolomic profile of relative intermediates normalized on protein content, of P160 *Tam-Cre;Pkd1^ΔC/flox^*cystic and control kidneys treated with ASOs. B. Quantification of Ki67 positive nuclei in the epithelium lining the cysts in kidney cortex of *Tam-Cre;Pkd1^ΔC/flox^* mice at P94, treated with *Scr-* or *Asns*-ASO. Data information: In A data are shown as mean ± SD. One-way ANOVA, corrected with Tukey’s multiple comparisons. In B data are shown as mean ± SD. Student’s unpaired two-tailed t-test. ns non- significant; *p<0.05; **p<0.01; ***p<0.001.

To further corroborate the metabolomic data highlighting the upregulation of the pyrimidine biosynthesis pathway in the long-term model of PKD, we investigated the expression of the first-step enzyme CAD in our animal models. In line with the increase in the products of the 3-step reaction of CAD (**Fig 5A**), we observed an increase in the activatory phosphorylation S1859 of the enzyme CAD in PBS- and *Scr*-treated kidneys at P160, which is partially reduced by *Asns* silencing (**Fig 6A**). Consistently, CAD was transcriptionally upregulated in different previously published datasets of PKD animal models, namely *Pkd1^v/v^* (**Fig 6B**; (Podrini *et al*, 2018)) and *Pkd1^RC/RC^* murine kidneys (**Fig 6C**; (Olson *et al*, 2019)), as well as in microarrays (**Fig 6D**; (Song *et al*, 2009)) and in snRNA-seq (**Fig 6E**; (Muto *et al*, 2021)) datasets from ADPKD patients, the last one mainly in the proximal tubule cluster of diseased kidneys. In addition, the analysis of previously published untargeted metabolomic data (Podrini *et al*, 2018) further confirmed an increase in the intermediates of *de novo* pyrimidine biosynthesis pathway downstream of CAD activity, in particular Dihydroorotate, Orotate and Orotidine in neonatal PKD kidneys from the *KspCre;Pkd1^ΔC/flox^* mouse model at P4 (**Fig 6E**).

**Figure 6.**
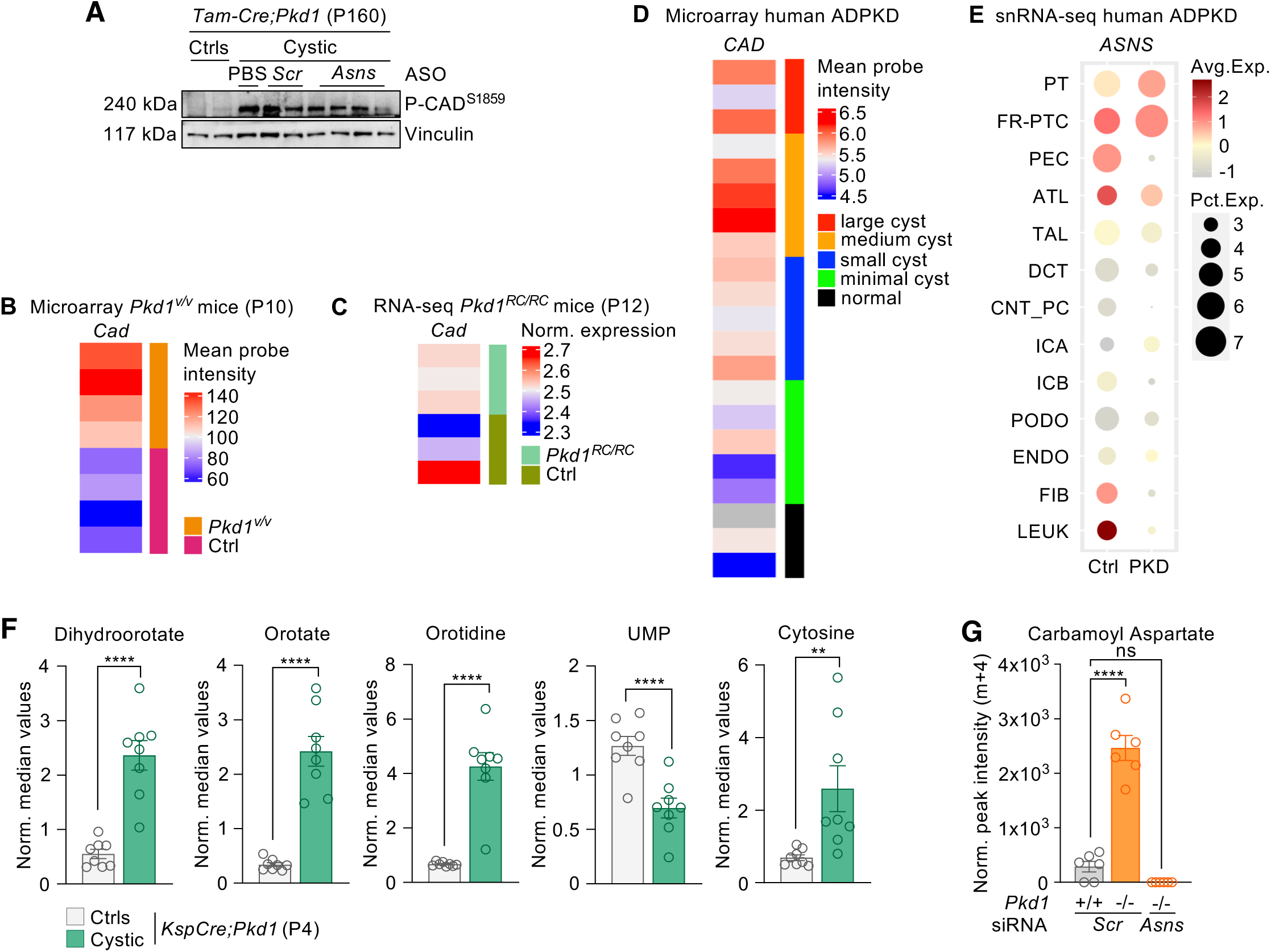
CAD-dependent pathway is upregulated in different models of PKD and rescued by *Asns* silencing *in vitro*. A. Western blot analysis of *Tam-Cre;Pkd1^ΔC/flox^* and relative controls at P160 treated with *Asns*- ASO or *Scr*-ASO. B. *Cad* expression in microarray from cystic kidneys of P10 *Pkd1^v/v^* mice, compared to relative controls. C. *Cad* expression in RNA-seq of *Pkd1^RC/RC^* cystic mice at P12, compared to controls. D. *CAD* expression in microarrays of ADPKD patients samples. E. Dot plot of snRNA-seq dataset showing *CAD* expression in clusters identified in human ADPKD cystic and normal kidney tissues. PT proximal tubule, FR-PTC failed-repair proximal tubular cells, PEC parietal epithelial cells, TAL thick ascending limb of Henle’s loop, DCT distal convoluted tubule, CNT_PC connecting tubule and principal cells, ICA Type A intercalated cells, ICB Type B intercalated cells, PODO podocytes, ENDO endothelial cells, FIB fibroblasts, LEUK leukocytes. F. Intermediate metabolites of *de novo* pyrimidine biosynthesis pathway analyzed through untargeted metabolomics of *KspCre;Pkd1^ΔC/flox^* cystic kidneys and relative controls at P4. G. Tracing metabolomics analysis (^13^C_5_-glutamine) evaluating labeled Carbamoyl Aspartate (m+4) in *Pkd1^-/-^* and control MEF cells ± silencing of *Asns*. Data information: in F-G data are shown as mean ± SEM. A-C Student’s t-test; D One-way ANOVA. ns not-significant; **p<0.01; ***p<0.001; ****p<0.0001.

Interestingly, glutamine is a key carbon and nitrogen source for the *de novo* pyrimidine biosynthesis. Based on the results above, we, therefore, assessed whether the CAD-dependent metabolic intermediates derive from *Asns*-driven glutamine utilization in *Pkd1^-/-^* cells. To this end, we used a ^13^C_5_-glutamine tracing experiment followed by LC-MS on *Pkd1^-/-^* MEF cells with or without silencing of *Asns* by siRNA (Podrini *et al*, 2018). We found an increase in labeled N-Carbamoyl aspartate (m+4) in *Pkd1^-/-^* cells. Importantly, the silencing of *Asns* in these cells completely restored the metabolite levels down to baseline (**Fig 6G**). Thus, we conclude that the required role of ASNS for glutamine utilization does not only fuel the TCA cycle and reductive carboxylation as we previously showed (Podrini *et al*, 2018), but it also strongly enhances the pathway of *de novo* pyrimidine biosynthesis likely explaining the dual role of ASNS in driving proliferation and allowing survival in cells lacking the *Pkd1* gene that we previously described.

### Analysis of non-rescued pathways identifies an opportunity for combination therapy in PKD

ASNS inhibition had a great effect on the metabolic derangement of PKD and rescued several pathways, and some of them to completion (i.e. the pathway was no longer significantly different from controls). However, we noticed that some other pathways were not as strongly affected (i.e. they were significantly improved over *Scr*-ASO, but still significantly different from controls). We thus set out to analyze the pathways for which we observed only a partial rescue by treatment with *Asns*-ASO, because in principle these should offer the opportunity to combine a second treatment to improve efficacy. Among these pathways, we found Glutathione metabolism, Nicotine and Nicotinate metabolism, and more pronouncedly glycolysis (**Fig 7A**). We previously proposed, based on our *in vitro* studies that glutamine compensates for the reduced utilization of glucose in the TCA cycle, and therefore combining glutamine with glucose starvation, or exposing cells silenced of *Asns* to glucose starvation further enhances the effective retardation of cell growth (Podrini *et al*, 2018). Thus, given that the glycolytic pathway was only partially affected in PKD kidneys treated with *Asns*-ASO prompted us to add a glycolytic inhibitor that we have previously characterized (2DG,(Chiaravalli *et al*, 2016)) to the *Asns*-ASO therapy. Therefore, we initiated a new study designed as the previous one (**Fig 2**), in which 2DG (2-deoxyglucose, the metabolically inactive analog of glucose) was added to the *Asns*-ASO or *Scr*-ASO treatment in the *Tam-Cre;Pkd1^ΔC/flox^* mouse model where cystogenesis was induced at P45 by Tamoxifen injection. Mice were next treated daily with gavage 2DG (100 mg/kg) or PBS and with either *Asns*-ASO or *Scr*-ASO (50 mg/kg), administered i.p. weekly up to P100 and every second week up to P160 (**Fig. 7B**). Results showed that the combined treatment reached a similar improvement of disease at the endpoint in *Asns*-ASO group as compared to *Scr*-ASO-treated mutants (P160, **Fig 7E** and **F**). However, combined treatment enhanced efficacy at an earlier time point as compared to *Asns*-ASO alone, as evidenced by a significant reduction in kidney volume measured by MRI scan at P100 (**Fig 7C** and **D**). Indeed, the side-by-side comparison between the two studies showed that all the animals enrolled in the combinatory study (COMBO) were consistently responsive to the treatment already at P100 in terms of kidney volume reduction, while in ASO study we observed a group of animals in which the response is appreciable only at later time points (**Fig 7G**). Collectively, these data indicate that adding an inhibitor of glycolysis (2DG) to the ASNS silencing enhances the effectiveness of the therapeutic approach at early time points, opening an opportunity for combination therapy by exploiting the reduced metabolic flexibility in PKD.

**Figure 7.**
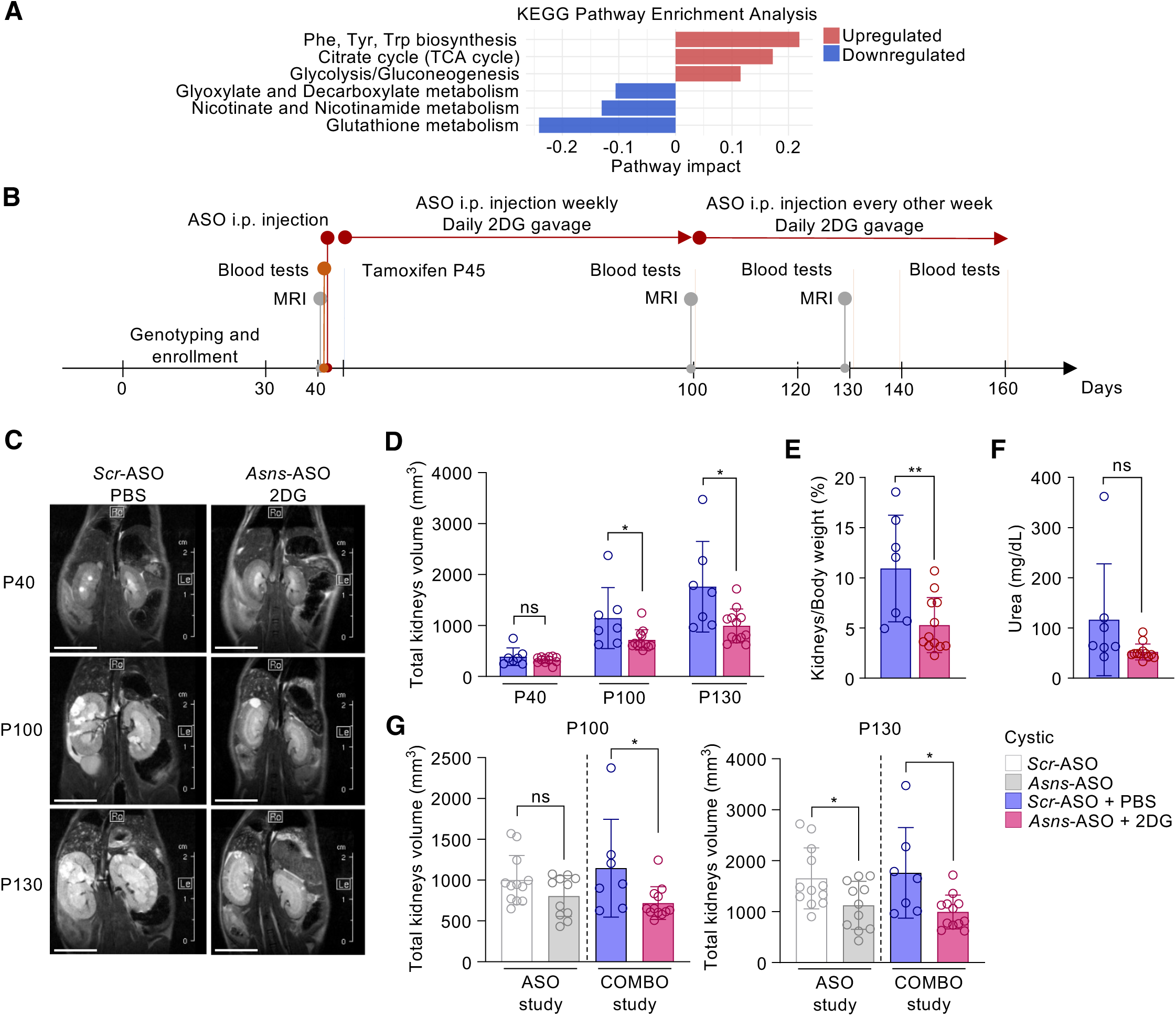
Combined targeting of glutamine usage via *Asns* and glycolysis delays the PKD progression in *Pkd1^ΔC/f^;TamCre* mice and ameliorates the end-point phenotype and function. A. Up and Down regulated Pathway Analysis Enrichment (cystic *Asns*-ASO not completely rescued compared to cystic *Scr*-ASO). B. Experimental design of *Tam-Cre;Pkd1^ΔC/flox^* and relative controls treated with *Asns*-ASO or *Scr*-ASO. C. MRI representative images of cystic kidneys treated with *Scr*-ASO+PBS or *Asns*-ASO+2DG. Images were acquired at P40 (before tamoxifen induction), P100 and P130. D. Total kidneys volume of cystic and control mice treated with *Scr*-ASO+PBS (n=7) or *Asns*- ASO+2DG (n=12), calculated at P40, P100 and P130. E. Percentage of kidneys weight normalized to total body weight of P160 mice treated with *Scr*- ASO+PBS (n=7) or *Asns*-ASO+2DG (n=12). F. BUN of cystic and control mice treated with *Scr*-ASO+PBS (n=7) or *Asns*-ASO+2DG (n=12), measured P160. G. Comparison of total kidneys volume measurement at P100 and P130 between cystic animals of the two presented studies: the single treatment with *Asns*-ASO (ASO study) and the combinatory treatment *Asns*-ASO + 2DG (COMBO study). ASO study bars correspond to data shown in Figure 2E, while COMBO study bars correspond to data shown in 7D. Data information: in A statistical significance of Pathway Enrichment and Impact was evaluated with Hypergeometric Test with Relative-betweeness Centrality, based on KEGG Database with p<0.05, FDR<0.1. In D-G data are shown as mean ± SD. Unpaired two-tailed Student’s t test. ns not- significant *p<0.05, **p<0.01.

## Discussion

In this study, we demonstrate that the metabolic enzyme ASNS is a valuable therapeutic target in Polycystic Kidney Disease. We show that this enzyme is upregulated in cells and tissues derived from murine or human PKD kidneys, and that using antisense-oligonucleotides (ASOs) directed against the murine *Asns* sequence results in a great amelioration of disease progression in orthologous PKD models (**Fig. 8**).

**Figure 8.**
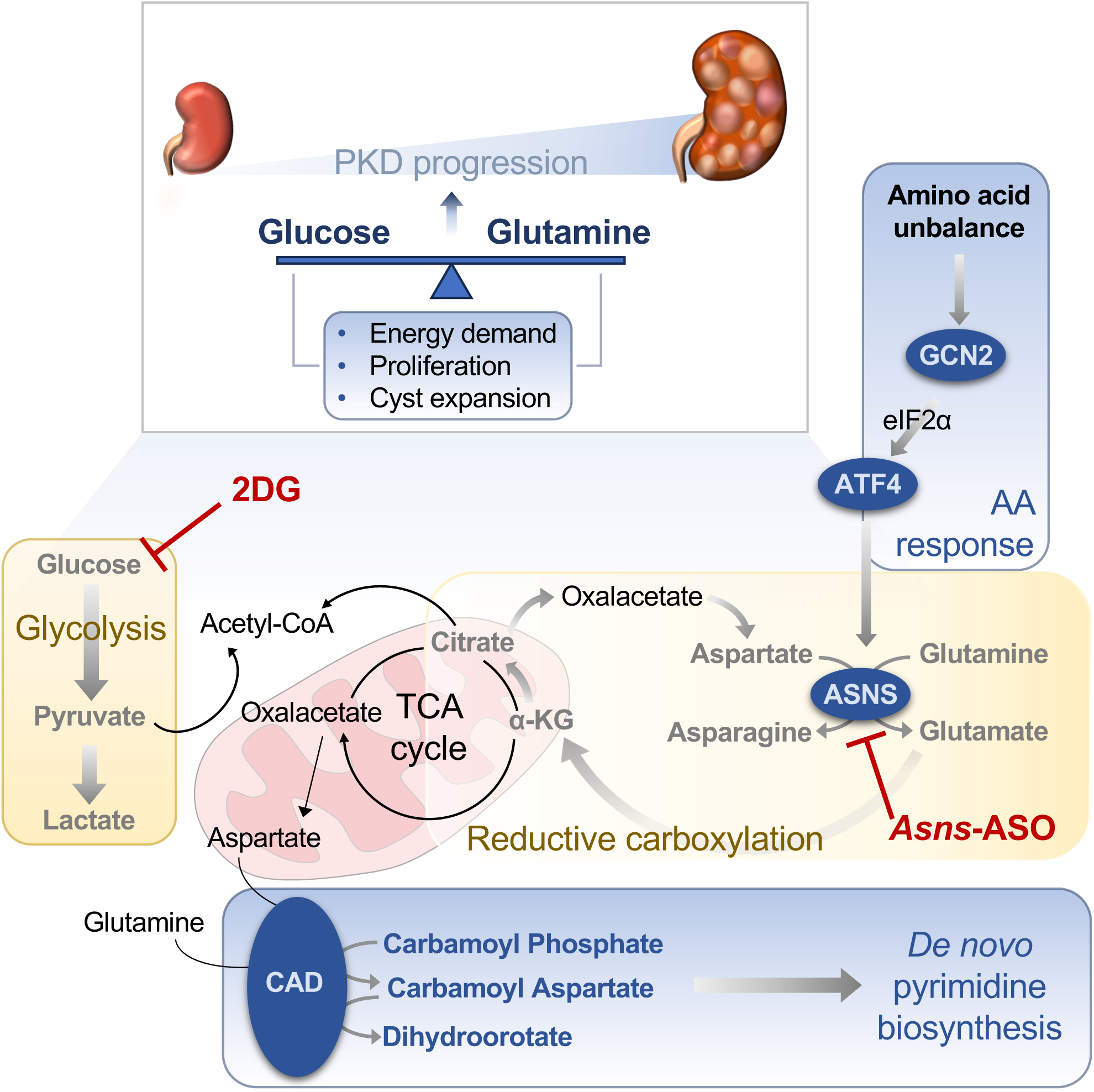
Schematic representation of metabolic rewiring supported by ASNS in PKD and proposed strategy for therapy. The scheme summarizes how glucose and glutamine usage support the progression of PKD. Increased glycolysis and lactate production (Rowe et al., 2013) reduce the fueling of glucose-derived carbons to the TCA cycle (Podrini et al, 2018) (yellow boxes). Glutamine utilization compensates for this deficit. Glutamine utilization is ASNS-dependent and fuels TCA cycle both through reductive carboxylation and oxidative phosphorylation (Podrini et al, 2018) (yellow boxes). Here we show that ASNS upregulation is downstream amino acid unbalance and GCN2-dependent AAR (blue box). In line with this, targeting ASNS *in vivo* in PKD models retards disease progression. Metabolomic profiling in these tissues reveals a prominent upregulation of *de novo* pyrimidine biosynthesis as a consequence of ASNS upregulation and glutamine utilization (blue box). Furthermore, combining targeting of ASNS and glycolysis via treatment with ASO and 2DG respectively breaks the energetic balance acquired in PKD resulting in the amelioration of cystic phenotype. Bold arrows and text highlight processes found upregulated in PKD.

In a previous study, we had demonstrated that glutamine utilization in PKD is compensatory for the reduced import of pyruvate into mitochondria owned to the enhanced lactate production resulting from a Warburg-like effect (Podrini *et al*, 2018). In addition, we had shown that this compensatory enhanced utilization of glutamine is driven by the enzyme asparagine synthetase (ASNS) a transamidase which removes an aminic group from glutamine to add it to aspartate, thus releasing asparagine and glutamate as products (Lomelino *et al*, 2017) (**Fig. 8**). In our previous studies, we had demonstrated that silencing ASNS *in vitro* completely prevents the utilization of glutamine in the TCA cycle, demonstrating that glutamate production for the TCA cycle depends on ASNS in cells mutant for the *Pkd1* gene (Podrini *et al*, 2018) (**Fig. 8**). Here, we provide evidence that ASNS is upregulated in cystic tissues *in vivo* as well, while expression of this enzyme in wild-type tissues is extremely low. Of great interest, and in line with the model we had proposed, a recent study found an increase in circulating asparagine in children and young adult PKD patients (Baliga *et al*, 2021), along with changes in the critical metabolites glutamine, glutamate and aspartate, products and substrates of the enzyme ASNS (Lomelino *et al*, 2017), providing strong supporting evidence of de-regulation of this pathway in ADPKD patients. In the current study we also show that changes in glutamine, glutamate, aspartate and asparagine can be observed in the cystic PKD kidneys, and corrected upon silencing of ASNS *in vivo*, demonstrating that the changes in levels of circulating metabolites in the ADPKD population are likely the result of their change in the kidneys, possibly also in patients. Indeed, we show that ASNS is upregulated in the kidneys of all the ADPKD mouse models and human ADPKD kidney dataset analyzed, as well as in all our *in vitro* and *in vivo* models of PKD, both at the level of the transcript and of the protein.

This extensive evidence of ASNS upregulation in murine and in patients’ tissues, along with our previous observation of the strong effect of *Asns* silencing in retarding growth in *Pkd1^-/-^* MEFs provided the rationale for targeting ASNS in an orthologous mouse model of ADPKD, which mimics the slow progression of the pathology observed in patients. We have designed and developed a set of anti-*Asns*-ASOs that strongly decrease the expression of this enzyme in cystic kidneys (collaboration with Ionis Pharmaceuticals^®^, see methods). Following in time the evolution of the main parameters characterizing the progression of PKD (i.e. increase in kidney volume and reduced renal function) revealed a quite striking decrease in kidney volume and weight, and a complete rescue of renal functionality down to the control levels upon administration of *Asns*-ASO as compared to controls (*Scr*-ASO). Of note, histological analysis of the *Asns*-ASO-treated kidneys revealed a quite variable phenotype not only between different animals, but also within each kidney with some samples presenting with portions of the kidneys that appeared completely rescued and others that appeared to remain somewhat cystic. This intra-sample variability was never observed in the controls and led us to propose that this effect is likely the result of a non-homogeneous distribution of the ASOs within the kidney. However, even when considering the great degree of variability of ASO distribution in the tissues, the improvement of disease manifestation and particularly of renal function was quite robust at the endpoint of the study. Of note, the study conducted in parallel on a group of controls treated with *Asns*-ASO revealed no indication of toxicity upon ASOs delivery, as assessed by body and kidney weight at sacrifice or overall appearance of the mice. These observations are relevant, because they show the potential to translate the approach of ASNS inhibition to humans. Indeed, the expression levels of this enzyme are quite low in adult tissues, with the only exception of the non-exocrine pancreas, explaining why the silencing of *Asns* seems to be well tolerated. Further studies in this sense should be conducted in the future. It should be noted here that targeting of ASNS has been proposed as a possible therapeutic approach, particularly in some solid tumors and in patients affected by acute lymphoblastic leukemia (Chiu *et al*, 2020). In this case, patients that tend to have high levels of circulating asparagine are subjected to treatment using asparaginase (ASNase) an enzyme able to degrade asparagine, which can be quite effective in these patients. However, some individuals develop resistance to the treatment due to an upregulation of the ASNS enzyme that can bypass the ASNase treatment by increasing the production of asparagine, thus leading to resistance to therapy (Gutierrez *et al*, 2006). It has been proposed that in principle such patients would also benefit from inhibition of ASNS. The mechanism behind the involvement of ASNS in PKD, however, might be slightly different, as we have previously demonstrated that ASNS is required intracellularly for the generation of glutamate which fuels the TCA cycle (Podrini *et al*, 2018). The role of asparagine seems to be less central in this disease, and in line with this observation we have found that treatment with ASNase does not improve (and possibly exacerbates) disease progression (Podrini and Boletta, unpublished). Thus, our data suggest that the production of glutamate (rather than that of asparagine) downstream of ASNS upregulation in PKD is central to disease progression and that the rationale for inhibiting ASNS might be quite different, though equally effective. One possible limitation of our studies stands in the fact that mice were treated with *Asns*-ASOs right from the beginning of disease initiation in the mouse, and it would be important to determine whether treatment at later stages of disease manifestation remains effective.

With respect to the mechanism leading to upregulation of ASNS, various previous studies have shown that while the protein is expressed at low levels in most tissues, it is strongly induced as a cellular response to cope with stress conditions, as part of activation of the integrated stress response (ISR).

The role of ISR, mainly the UPR branch, in the context of cystic kidney disorders is still debated. Indeed, it has been described that activation of the PERK-dependent branch and activation of the downstream ATF4 transcriptional program contributes to establishment of juvenile cystic kidney ciliopathy (Panda *et al*, 2022). On the other hand, Fedeles et al. demonstrated that activation of the UPR pathway plays a protective role against cyst formation, but that the pathway itself is not deregulated in those kidneys (Fedeles *et al*, 2015). Here, we demonstrated that the GCN2-dependent AAR branch of the ISR is strongly activated in ADPKD models, and that it drives the transcription of ASNS, since silencing of GCN2 in *Pkd1^-/-^*cells is sufficient to restore lower levels of ATF4 and ASNS (**Fig. 8**). Thus, our data imply that an alternative route of activation of the pathway, dependent on amino acid deregulation, can drive this response in PKD, in line with previous studies showing amino acids alteration in PKD (Ramalingam *et al*, 2021). But why is ASNS so essential for cyst growth and what are the consequences of its upregulation? Metabolomic profiling of ASO-treated kidneys sheds light on how ASNS can orchestrate multiple metabolic processes to support the progression of the pathology, and also provide evidence of how treatment could impact on this specific aspect. Despite the fact that we found several metabolic pathways de-regulated in the PKD kidneys at P160 and that most of these are rescued by ASOs treatment, it should be noted that the strength of rescue can be very different from pathway to pathway, with some being completely restored back to controls and other being less positively affected by *Asns*-ASO. There could be multiple explanations for these findings. One possibility is that the pathways that are fully rescued are more closely related to the enzyme ASNS and to glutamine utilization and therefore are more robustly impacted. The second possibility is that some pathways have a lower flexibility than others and cannot be compensated by carbon sources other than glutamine in PKD. In all cases, the non-homogeneous rescue of metabolic pathways seems to suggest that the metabolic rescue is not merely secondary to rescue of disease progression, as in this case one would expect more homogeneous changes. Indeed, we think that this observation demonstrates that the ASO treatment effect on metabolic processes cannot be only reconducted to phenotypic rescue and restoration of healthy tissue. Importantly, the metabolomic profile corroborated the mechanism that supports the upregulation of ASNS in PKD. Indeed, we defined a profound unbalance in amino acid levels in PKD kidneys, which supports the transcriptional regulation of the enzyme via GCN2-dependent AAR activation. The metabolomic profiling also provided important information on how ASNS targeting retards PKD progression. In a previous study we showed that PKD cells *in vitro* are more sensitive to reduced proliferation upon *Asns* silencing as compared to controls (Podrini *et al*, 2018). Here, we demonstrated that ASO treatment almost completely rescued the upregulation of *de novo* pyrimidine biosynthesis in diseased kidneys, thus resulting in reduced proliferation and consequent cyst formation. Moreover, we defined a central role for ASNS in the proficient usage of glutamine in PKD cells, which can be used to sustain the CAD-dependent step of nucleotide production, essential for proliferation (**Fig. 8**). This model is also well supported by the results obtained *in vivo*.

Among the metabolic pathways that remain only partially corrected by *Asns*-ASO is that of glycolysis in line with the idea that the ASNS-mediated utilization of glutamine provides the carbon source for TCA cycle feeding, in conditions in which enhanced glycolysis depletes pyruvate import to mitochondria, but it is not the main provider of energy in these cells, which is almost entirely driven by glucose (Rowe *et al*, 2013; Podrini *et al*, 2018). Thus, given the effectiveness of the inhibitor of glycolysis 2-DG in improving PKD progression reported by multiple groups including ours (Chiaravalli *et al*, 2016), we tested whether combining the two strategies of interference with glucose utilization (by 2DG) and of glutamine utilization (by *Asns*-ASO) would act synergistically and provide the additional benefit that we would predict based on our *in vitro* data (Podrini *et al*, 2018). Indeed, we show that 2DG provides an additional improvement of disease amelioration *in vivo* over treatment with *Asns*-ASOs alone. The effect is robust, but does not completely rescue disease progression, suggesting that additional ways of escape are available to the diseased PKD mutant cells. One important question that remains unanswered is how the multiple metabolic alterations observed in ADPKD originate and what is the connection with the Polycystins function. Recent studies have described that the cleaved C-tail of PC-1 translocates to mitochondria and interacts with multiple metabolic enzymes (Lin *et al*, 2018; Onuchic *et al*, 2023; Lin *et al*, 2023; Pellegrini *et al*, 2023), although the molecular details of how these multiple interactions might result in the observed metabolic phenotype remain currently unexplained.

In this context, it should not be ignored that the polycystins are ciliary proteins. In a recent study, we demonstrated that primary cilia respond to glutamine levels and that they do so via the enzyme ASNS, which localizes at the base of cilia (Steidl *et al*, 2023). In that case, we have shown that removal of primary cilia from cells results in a metabolic reprogramming with respect to glucose and glutamine utilization, that is the opposite as compared to the one that we have observed in ADPKD, i.e. cells utilize reduced levels of glucose and reduced levels of glutamine. These data suggest that metabolic reprogramming might be part of the CDCA pathway, described to be a cascade constitutively activated at cilia upon removal of the polycystins (Ma *et al*, 2013). While the molecular nature of this cascade remains obscure, the existence of such a pathway seems to be demonstrated by the fact that removal of cilia unexpectedly improves disease progression in the context of *Pkd1* or *Pkd2* mutants (Ma *et al*, 2013). Here, we propose that ASNS might be a key component of this CDCA pathway. Indeed, this enzyme seems to respond to all criteria for a molecular definition of the CDCA pathway: i) it is upregulated in the absence of the polycystins (Podrini *et al*, 2018 and current work), but downregulated in the absence of cilia (Steidl *et al*, 2023); ii) it localizes at primary cilia (Steidl *et al*, 2023); iii) its inhibition results in renal cysts improvement (Current work). Further studies should be undertaken to better understand this aspect, while the main conclusion of our current work is that we have identified a novel and quite robust new target for therapy in ADPKD, the enzyme ASNS.

## Materials and Methods

### Cell culture and treatment

Immortalized *Pkd1^+/+^* and *Pkd1^−/−^* MEFs (Distefano et al, 2009 *Mol. Cell Biol.*) were cultured in DMEM (Gibco, #5030), supplemented with 10% FBS (Thermo Fisher, #26400-044), 44 mM sodium bicarbonate (Sigma Aldrich, #S-6014), 1% Pen/Strep (Gibco, #15070-63), 4 mM glutamine (Gibco, #25030) 25 mM glucose (Sigma Aldrich, #G5767). For glucose deprivation, cells were seeded in complete medium, and subjected to glucose deprivation for 24h.

### siRNA

For transient silencing of *Asns* (Ambion, #AM16704/188316) and *Eif2ak4*, 20 nM predesigned siRNA (Ambion, #AM16704/4390815) have been used along with scrambled control, following the manufacturer’s instructions. 100 000 cells/well have been seeded in 6-wells plate for siRNA transfection, in DMEM 10% FBS medium, without antibiotics. Transfection has been performed for 2 consecutive days, reaching a final concentration of 30 nM siRNA, using Lipofectamine RNAiMAX (ThermoFisher, #13778150), following the manufacturer’s instructions. Cells have been processed for RNA and protein extraction after 72 hours from the first transfection.

### Generation of Tam-Cre;Pkd1^ΔC/flox^ and KspCre;Pkd1^ΔC/flox^ mice

C57/BL6 *Tam-Cre;Pkd1^ΔC/+^* were crossed with C57/BL6N *Pkd1^flox/flox^* to generate the *Tam- Cre;Pkd1^ΔC/flox^*mice (Chiaravalli *et al*, 2016). Cre recombinase activity was induced by a single injection of Tamoxifen (250 mg/kg; Sigma Aldrich, #T5648) at P45 or P25 for a long-term model involved in *Asns*-ASO study and the previous pilot study, respectively. Tamoxifen was freshly prepared and dissolved in corn oil (#C8267; Sigma Aldrich) by continuous shaking at 37°C for at least 6 hours and was injected intraperitoneally. For the generation of *KspCre;Pkd1^ΔC/flox^*we crossed C57/BL6N *Pkd1^flox/flox^* with *KspCre;Pkd1^ΔC/+^*, as previously described (Rowe *et al*, 2013). Animal care and experimental protocols were approved by the Institutional Care and Use Ethical Committee at the San Raffaele Scientific Institute and further approved by the Italian Ministry of Health (IACUC 736 and IACUC 921).

### Treatment with ASOs *in vivo*

*Tam-Cre;Pkd1^ΔC/flox^* mice and relative controls were randomly distributed in the group of treatment for each study. Detection of increased kidney volume or presence of cysts in kidneys and/or in the liver during the first MRI scan represented exclusion criteria for the study. After enrolment, mice were treated with antisense oligonucleotide targeting *Asns* (*Asns*-ASO; TATTTTATCACACTCC) or non-targeting scramble control (*Scr*-ASO; ACGATAACGGTCAGTA) (Ionis Pharmaceuticals^®^). As the efficacy/tolerability was dosed at 50 mg/kg/week, ASOs were administered at the same dosage via weekly intraperitoneal injection for the first 2 months after Tamoxifen induction, and every second week after P100 till the end of the experiment. In the pilot experiment, ASOs were administered weekly at the same dosage till P60 and every second week until the end of the study. For the combinatory approach, experimental animals were treated daily (5 days per week) with 100 mg/kg 2DG (Sigma Aldrich, #D8375) gavage (combined with *Asns*-ASO treatment) or PBS gavage (combined with *Scr*-ASO treatment).

During all the studies presented, animals were subjected to 5 blood tests (P40, P100, P130, P140, P160) to monitor renal and liver parameters and 3 MRI scans (P40, P100, P130) to monitor kidney volume.

At the end of the study, animals were weighed and perfused with cold PBS after anesthesia. Kidneys were collected and weighed to calculate kidneys/body weight. Tissues were then processed for *ex-vivo* analysis.

### Biochemical serum analysis

Urea (#0018255440) and Crea (#0018255540) were used for the quantitative determination of the serum level with an International Federation of Clinical Chemistry and Laboratory Medicine optimized kinetic ultraviolet (UV) method in an ILab650 chemical analyzer (Instrumentation Laboratory). Urea and Crea are expressed as mg/dl. SeraChem Control Level 1 and Level 2 (#0018162412 and #0018162512) were analyzed as quality control.

### MRI

All MRI studies were performed on a 7-T Preclinical Scanner (BioSpec 70/30 USR, Paravision 6.0.1; Bruker) equipped with 450/675 mT/m gradients (slew rate: 3400–4500 T/m per second; rise time: 140 µs) and a circular polarized mouse body volume coil with an inner diameter of 40 mm. Mice were under general anesthesia obtained by 1.5%–2% isoflurane vaporized in 100% oxygen (flow of 1 L/min). Breathing and body temperature were monitored during MRI (SA Instruments, Inc., Stony Brook, NY) and maintained around 30 breaths per minute and 37°C, respectively. All MRI studies included axial and coronal RARE T2–weighted sequences (slice thickness, 0.6 mm; interslice gap, 0 mm) with several slices, allowing complete coverage of both kidneys. Manual segmentation of the kidneys was performed on each slice using NIH software MIPAV (version 7.4.0), excluding the collecting system, and kidney volume results from automatic summation of voxel volumes. Both acquisition and analysis of the data were performed blindly by an operator unaware of genotype or treatment conditions.

### Histology and immunohistochemistry

After euthanasia, kidneys were collected, washed in PBS, weighed, and fixed in 10% neutral buffered formalin containing 4% formaldehyde (Bio-Optica, 05-01V15P). Tissues were transferred to 70% ethanol after 24 hours, and paraffin embedded for subsequent analysis. Formalin-fixed paraffin-embedded consecutive sections (4 um) were dewaxed and hydrated through graded decrease alcohol series and stained for histology or immunohistochemical characterization (IHC).

For histological analysis in bright-field microscopy, slides were stained using standard protocols for Hematoxylin and Eosin (using Mayer’s Hematoxylin, BioOptica #05-06002/L and Eosin, BioOptica #05-10002/L).

For IHC characterizations, slides were immunostained with Automatic Leica BOND RX system (Leica Microsystems GmbH, Wetzlar, Germany). First, tissues were deparaffinized and pre-treated with the Epitope Retrieval Solution (ER1 Citrate Buffer) at 100°C. Primary antibody against Ki-67 ((D3B5) Rabbit mAb (Mouse Preferred; IHC Formulated) (CST, #12202)) was used 1:200 and developed with Bond Polymer Refine Detection (Leica, DS9800).

Slides were acquired with Aperio AT2 digital scanner at a magnification of 20X (Leica Biosystems) and analyzed with Imagescope (Leica Biosystem).

### Cystic index

Images of transversal sections from the inner part of the kidney after Hematoxylin-Eosin staining were taken (Zeiss AxioImager M2m with AxioCam MRc5) at 2.5X magnification. The ImageJ program (http://rsb.info.nih.gov/ij/) was then used to quantify the total surface of the kidney and the total cystic area. We next calculated the ratio between the cystic area and the total area of the kidney.

### Ki67 quantification

Proliferative Ki-67-positive cells have been quantified semi-automatically. Ki-67 nuclear staining have been automatically detected in the epithelium lining cortical cysts by Aperio Image analysis software (Leica Biosystems). Positive nuclei have been manually counted and normalized on the total number of nuclei in cyst epithelium.

### Ultra-High-Pressure Liquid Chromatography-Mass Spectrometry (MS) metabolomics experiments

Kidney tissues were grinded in dry ice and weighed. 15 mg of powder was extracted in 1 mL of ice-cold extraction solution (methanol:acetonitrile:water 5:3:2 v/v/v) (Nemkov *et al*, 2017). Suspensions were vortexed continuously for 30 min at 4°C. Insoluble material was removed by centrifugation at 18,000 g for 10 min at 4°C and supernatants were isolated for metabolomics analysis by UHPLC-MS. Analyses were performed using a Vanquish UHPLC coupled online to a Q Exactive mass spectrometer (Thermo Fisher, Bremen, Germany). Ten microliters of sample extracts were loaded onto a Kinetex XB-C18 column (150x2.1 mm i.d., 1.7µm – Phenomenex). Samples were analyzed using a 5 minutes method as described (Nemkov *et al*, 2017; Reisz *et al*, 2019).

Metabolite assignments were performed using MAVEN (Princeton, NJ, USA) (Clasquin *et al*, 2012). Metabolites levels were then normalized to BCA protein quantification (Pierce™ BCA Protein Assay Kits 23225). Graphs and statistical analyses (either PCA, HCA or MetPA) were prepared with GraphPad Prism 10.0 (GraphPad Software, Inc, La Jolla, CA), GENE-E (Broad Institute, Cambridge, MA, USA), MetaboAnalyst 5.0 (Chong *et al*, 2018), and RStudio Team (2020). RStudio: Integrated Development for R. RStudio, PBC, Boston, MA URL http://www.rstudio.com/.

### Ultra-High-Pressure Liquid Chromatography-Mass Spectrometry (MS) tracing metabolomics experiments

Tracing metabolomics experiment on *Pkd1^-/-^* and control MEF cells has been conducted as previously described (Podrini *et al*, 2018). Briefly, cells were cultured in either culture media enriched with ^13^C_5_ Glutamine (Cambridge Isotope Laboratories, CLM-1822-H-PK). Metabolite assignments and ^13^C_5_ Glutamine tracing experiments, isotopologue distributions, and correction for expected natural abundances of 13C isotopes, were performed using RStudio AccuCor (Su *et al*, 2017). Metabolites levels were then normalized to BCA protein quantification (Pierce™ BCA Protein Assay Kits 23225).

### Real-time PCR Analysis

RNA was isolated from plated cells or snap-frozen kidneys using the RNAspin Mini kit (GE Healthcare, #25-0500-72) according to the manufacturer’s instructions. For reverse transcription of RNA, Oligo(dT)15 primers (Promega, #c1101) and ImProm-II Reverse Transcriptase (Promega, #A3802) were used. Quantitative real-time PCR analysis was performed on duplicates using SYBR Green I master mix (Bio-Rad, 48873) on CFX96 Touch Real-Time PCR Instrument (Bio-Rad). Primers sequences:

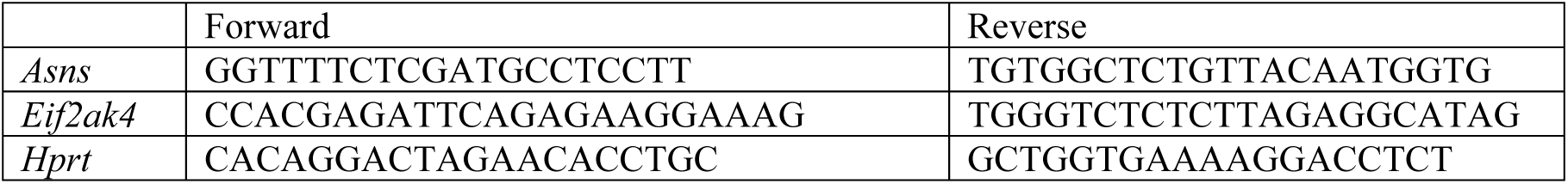

### Western Blot Analysis

Kidneys were grinded on dry ice and re-suspended in lysis buffer (10 mM Hepes, 10 mM KCl, 1.5 mM MgCl_2_, 0.34 M sucrose, 1 mM dithiothreitol, 10% glycerol [pH 7.9], 0.1% Triton X-100, complete protease inhibitors (Roche, #4693132001), and phosphatase inhibitors (1 mM final concentration of glycerophosphate, sodium orthovanadate, and sodium fluoride). After centrifugation, we separated the supernatant and the pellet. The supernatant was clarified by a second centrifugation. Next, we used 5% skim milk in Tris-buffered saline and Tween-20 (Sigma Aldrich, P1379) for blocking. Primary antibodies were diluted in 1x TBS-T supplemented with 3% of BSA (Sigma Aldrich, #A7906). HRP-conjugated secondary antibodies were diluted 1:10.000 (anti-Mouse, Life

Technologies, #A11003; anti-Rabbit, Life Technologies, #11035) (or more if necessary) in 5% of skim milk, 1x TBS-T, and detected by ECL (Cytiva, #RPN2106) alone or supplied with 10% of SuperSignal West Femto (Thermo Fisher Scientific, #34095) when required.

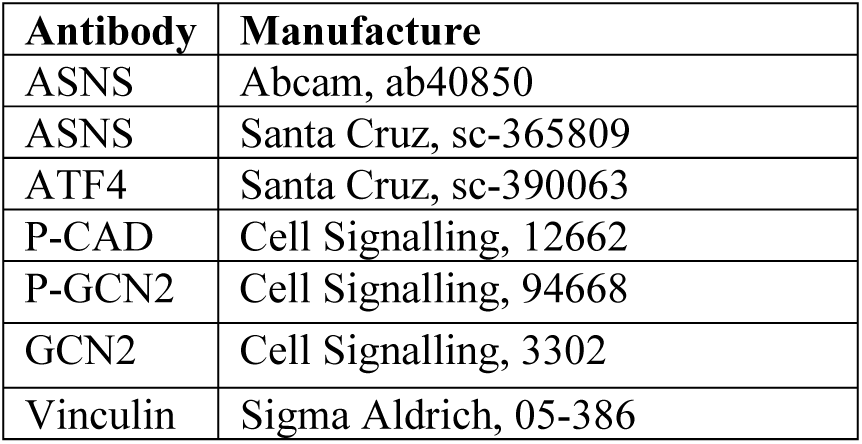

| Antibody | Manufacture |
| --- | --- |
| ASNS | Abcam, ab40850 |
| ASNS | Santa Cruz, sc-365809 |
| ATF4 | Santa Cruz, sc-390063 |
| P-CAD | Cell Signalling, 12662 |
| P-GCN2 | Cell Signalling, 94668 |
| GCN2 | Cell Signalling, 3302 |
| Vinculin | Sigma Aldrich, 05-386 |

### Datasets sources

Normalized counts were obtained directly from publications (Olson *et al*, 2019), normalized probe intensity data were extracted from the series matrix files from the GEO repository with accession numbers: GSE7869 (Song *et al*, 2009) and GSE121563 (Podrini *et al*, 2018). Microarray probes of each data set were aggregated on a gene basis by averaging normalized probe intensities.

### Data Visualization

scRNA-seq dotplots visualizing the expression of controls and ADPKD patients were created using the “Kidney Interactive Transcriptomics” platform http://humphreyslab.com/SingleCell (Wu *et al*, 2022). Expression heatmaps of RNA-seq and Microarray expression data were created using R (R Core Team 2022, URL https://www.R-project.org/) and the ComplexHeatmap package (Gu *et al*, 2016).

### Statistical analysis

Statistical analyses were performed using Prism 10, GraphPad Software. *t*-test was used for all analyses comparing two groups. ANOVA statistical analysis followed by Tukey’s multiple comparison test was performed in all analyses where more than two groups were present.

## ACKNOWLEDGEMENTS

The authors are grateful to the other members of the Boletta laboratory for useful discussions. This work was supported by the Italian Ministry of Health (RF-2018-12368254 to AB; GR-2016-02364851 to CP), the Italian association of patients with PKD (AIRP to AB), by the Italian association for research on cancer (AIRC, IG2019-23513 to AB). The authors are grateful to Dr. Silvia Bramani for her continuous support.

## AUTHOR CONTRIBUTIONS

C.P. conceived the project, performed the experiments and analysis, prepared some of the figures. S.C. Performed experiments, performed analysis, conceived and performed part of the studies, wrote the manuscript, prepared the figures; L.T. and D.S performed analysis of metabolomic data, prepared graphs; T.C. performed MRI imaging and volumes calculation; M.C. performed in vivo experiments, analysed data, helped designing the experiments; D.S. performed analysis of transcriptional profile datasets; S.C. performed tracing metabolomic experiments; A.S. supervised the animal imaging experiments and volume evaluations; A.D’A. supervised and performed metabolomic analysis; C.F supervised and discussed metabolic tracing data; A.B. supervised some of the metabolomics studies and contributed discussion and data revision; A.B. Conceived the studies, supervised the work and collaborations, wrote the manuscript.

## The paper explained

### Problem

Autosomal Dominant Polycystic Kidney Disease (ADPKD) is one of the most common monogenic disorders affecting humans. The disease manifests with the formation of cysts in both kidneys. Cysts are enclosed structures of epithelia that grow over time leading to the compression of the surrounding normal parenchyma, eventually causing loss of renal function. Only one compound, Tolvaptan, able to slow down growth and retard progression is currently available to patients, but it presents with limited efficacy and limited tolerability. We and others described a peculiar role of metabolic reprogramming in PKD which favors growth of the cysts and have proposed that this cellular dysfunction represents an important vulnerability to be targeted for therapy.

### Results

Here we used antisense oligonucleotides (ASOs) to reduce the expression levels of the metabolic enzyme asparagine synthetase (ASNS) in orthologous and slowly progressive ADPKD murine models. We report that treatment with such ASOs greatly improves disease progression and restores almost normal function in mice. The effect of silencing prominently improves the metabolic dysfunction in PKD tissues and identifies novel metabolic vulnerabilities. We also show that combining a glycolysis inhibitor (2DG) with silencing of ASNS further ameliorates disease manifestation.

### Impact

We have identified a novel target for therapy in Polycystic Kidney Disease, whose targeting results in great amelioration of disease progression and regression of several metabolic alterations. The results might lead to the development of a much needed novel therapy for this disorder.

## Expanded View Figure legends

**Figure EV1**

A. Experimental design of pilot study on *Tam-Cre;Pkd1^ΔC/flox^*and relative controls treated with *Asns*-ASO or *Scr*-ASO. (n=4 ctrl *Scr*-ASO; n=2 ctrl *Asns*-ASO; n=4 cystic *Scr*-ASO; n=4 cystic *Asns*-ASO)

B. *Asns* mRNA expression in *Tam-Cre;Pkd1^ΔC/flox^* and relative controls treated with *Asns*-ASO or *Scr*-ASO

C. Representative images of cystic kidneys and relative controls at P94 treated with *Asns*-ASO or *Scr*-ASO

D. Percentage of kidneys weight normalized to body weight of cystic and relative controls treated with *Asns*-ASO or *Scr*-ASO.

E. BUN of cystic and relative control kidneys treated with *Asns*-ASO or *Scr*-ASO.

F. Quantification of the cystic area percentage of the total kidney area measured in transversial sections of cystic *Scr*-ASO or *Asns*-ASO groups.

Data information: in B, D, E data are shown as mean ± SD. One-way ANOVA. ns non-significant; *p<0.05; **p<0.01. In F data are shown as mean ± SD. Student’s unpaired two-tailed t-test. **p<0.01

**Figure EV2**

A. Dedrogram diagram showing the hierachical clustering of 4 ASO-treated groups of samples analyzed through LC-MS.

B. Heatmap based on the HCA of the metabolom (265 metabolites) comparing *Scr-* and *Asns*- ASO treated cystic and control kidneys.

C. Heatmap based on the HCA of amino acids detected in PKD cystic and control kidneys treated with *Scr*-ASO or *Asns*-ASO.

Data information: in A-C clustering based on t-test/ANOVA result performed with Metaboanalyst 5.0.

